# Viral Sentry AI (VirSentAI) - Automated Zoonotic Surveillance & Drug Repurposing Agent

**DOI:** 10.64898/2025.12.29.684576

**Authors:** Cristian R. Munteanu, José M. Vázquez-Naya, Eduardo Tejera

**Author notes:** Corresponding author: Prof. Dr. Cristian R. Munteanu.

## Abstract

Zoonotic viruses capable of jumping from animal reservoirs into human populations represent a persistent and unpredictable menace to global health. To confront this challenge, we developed Viral Sentry AI (VirSentAI), an autonomous agent designed to close the gap between viral emergence and therapeutic response. Unlike static analysis tools, VirSentAI operates as a continuous sentinel, automatically scanning public databases (e.g., NCBI) for new viral genomes and executing a three-stage agentic surveillance workflow, with distinct, specialized AI architectures for generated text, macromolecule sequences, and drug chemical data. First, the system is using gemma-2-9b, utilizing this Large Language Model to parse unstructured submission records and extract critical meta-information that provides context to the raw data. In the second stage, the system employs a novel deep-learning topology, virsentai-v2-hyena-dna-16k, a fine-tuned HyenaDNA model capable of processing complete viral genomes up to 160,000 bases. This architecture captures subtle, long-range genomic dependencies to predict human infectivity with high precision.

Upon flagging a high-risk pathogen, the agent autonomously triggers a downstream therapeutic module as the stage three. It extracts NCBI viral protein sequences and utilizes a PLAPT (Protein-Ligand Affinity Prediction Transformer) model to calculate affinity interactions against the ChEMBL-curated set of approved therapeutics, instantly identifying candidates for drug repurposing. In rigorous cross-validation on a curated dataset of 31,728 complete viral genomes, the surveillance module demonstrated robust discriminatory power, achieving an AUROC of 0.95 in classifying human host potential. By integrating state-of-the-art genomic modeling with automated lead compound screening, VirSentAI offers a proactive, end-to-end solution for pandemic preparedness. The platform is freely accessible at https://muntisa.github.io/virsentai, with source code available at https://github.com/muntisa/virsentai.

## 1. Introduction

The prediction of viral spillover from animal reservoirs to humans represents a critical frontier in global public health security. The societal disruption caused by the COVID-19 pandemic served as a stark reminder that most emerging human pathogens, including coronaviruses, filoviruses, and influenza viruses, are of zoonotic origin. In principle, genomic sequencing of a novel animal virus could determine its potential to infect humans. In practice, however, laboratory-based host-range determination is a resource-intensive and retrospective process. This limitation has catalyzed the development of computational models designed to rapidly assess zoonotic risk from viral sequence data alone, sifting through vast genomic archives to identify potential threats.

The landscape of these predictive models reveals a clear evolutionary trajectory. Initial efforts produced powerful, yet highly specialized, tools for specific viral families. For instance, HostPredictor, a gradient boosting ensemble leveraging codon usage bias and k-mer frequencies, achieved a high area under the receiver operating characteristic curve (AUC) of 0.95 for avian influenza viruses (Brierley, Mould-Quevedo, & Baylis, 2024). Similarly, Flu-CNN, a deep 1D convolutional neural network, demonstrated 99% accuracy in identifying the host specificity of Influenza A viruses (Hu et al., 2025). While potent, the utility of such models is intrinsically narrowed to well-defined genomic contexts.

Subsequently, broader strategies emerged to capture more universal signatures of zoonotic potential. Foundational work by Babayan’s team (Babayan, Orton, & Streicker, 2018) demonstrated that evolutionary signatures within viral genomes could effectively predict reservoir hosts, stimulating numerous subsequent investigations. This line of inquiry led to tools like VIDHOP, which applied deep neural networks to partial genomic sequences from diverse viral families, achieving high AUCs between 0.93 and 0.98 for specific viruses like Rabies and Rotavirus (Gong, Qian, & Poon, 2021). Another approach, BiLSTM-VHP, utilized a bidirectional LSTM network to analyze patterns in short viral genome fragments (Efat, Islam, Rasel, & Haque, 2025). A persistent tension exists in these approaches, balancing the precision of family-specific models against the broader applicability of pan-viral systems, which can sometimes sacrifice contextual depth. As highlighted in a recent review, model performance is highly context-dependent, with no single tool demonstrating universal optimality (Kim et al., 2025). This fragmentation necessitates that researchers often navigate a complex ecosystem of tools, each with distinct data requirements and limitations.

The advent of large-scale, foundational language models trained on genomic data has introduced a paradigm shift. Evo 2, a scale-unifying model pretrained on 9.3 trillion base pairs from across all domains of life, exemplifies this new frontier (Merchant, King, Nguyen, & Hie, 2025). Its StripedHyena 2 architecture can process sequence contexts up to one million nucleotides, enabling the recognition of long-range dependencies invisible to previous models. By encoding a deep representation of genomic grammar, Evo 2 provides latent features richly attuned to evolutionary constraints and regulatory motifs—attributes highly relevant for host-prediction tasks.

A significant constraint, however, is that Evo 2 was not trained on known human-pathogen viral genomes, an intentional precaution against misuse (Merchant et al., 2025). This exclusion prevents its direct application for predicting human infectivity without further fine-tuning. Nonetheless, the existence of such foundational models reshapes the development landscape.

VirSentAI was developed to occupy this precise niche as a three AI model agent processing text, DNA and protein sequences, and drug SMILES. It is an AI-driven web service integrating a modern, large-scale sequence model with an automated surveillance platform. By fine-tuning a HyenaDNA architecture, VirSentAI leverages the power of large-scale pre-training to scan entire viral genomes, flag those with the highest likelihood of infecting humans. PLAPT pre-trained model was used to propose automatic drug repurposing for the viral proteins. The host virus was corrected with a pre-trained LLM such as MedGemma using other fields. VirSentAI is designed for accessibility, operating as an autonomous system that periodically scans public repositories like NCBI for new viral sequences and presents risk assessments and drug repurposing solutions without requiring bioinformatics expertise from the user. VirSentAI aims to streamline zoonosis surveillance by making an advanced, broadly applicable host-prediction model accessible to the wider virology and epidemiology community.

## 2. Methods

Our tool’s foundation rests on a blend of genomic insight and computational finesse. We subdivided the build to keep things clear. Behind the scenes, VirSentAI runs a set of automated scripts that routinely comb through the NCBI database, searching for newly released complete viral DNA sequences. These sequences are then passed into our updated prediction engine— virsentai-v2-hyenadna-16k—which estimates the likelihood that each virus could infect humans. The results are not just numbers on a screen; they are translated into ranked tables and clear visual summaries, highlighting the viruses that deserve a closer look. **Figure 1** shows the conceptual workflow of the VirSentAI platform, from the initial input of a viral sequence to the final output of a zoonotic risk score and drug repurposing for viral proteins, all integrated within an automated surveillance pipeline.

**Figure 1:**
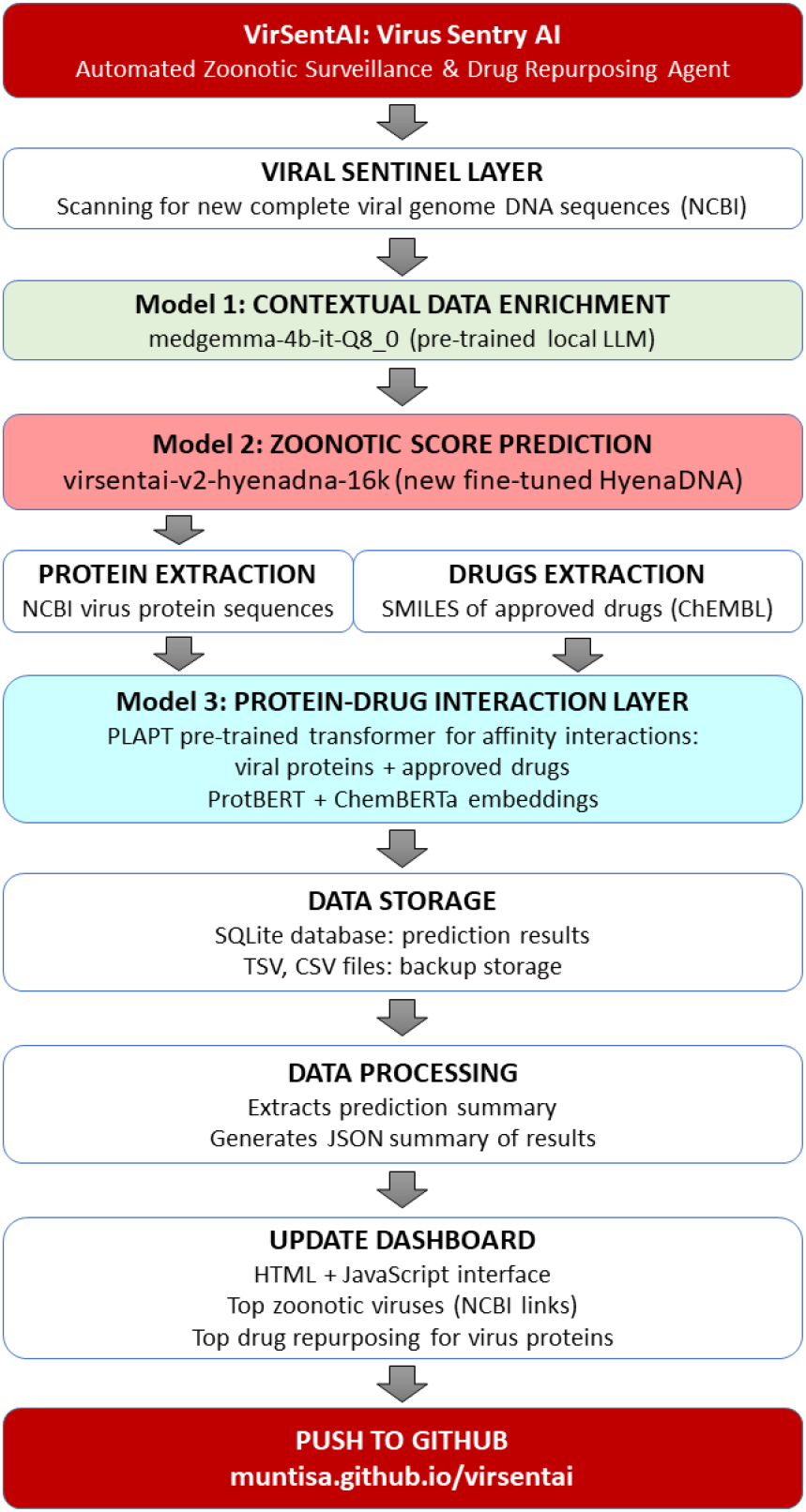
VirSentAI platform overview diagram illustrating the flow from input genome to the virsentai-v2-hyenadna-16k model, the resulting zoonotic risk score and automatic drug repurposing solution.

VirSentAI is a multimodal agent for zoonotic defense & therapeutic data fusion, an autonomous, tri-model agent designed to close the gap between viral emergence and therapeutic response. It orchestrates a unified surveillance workflow by continuously synthesizing intelligence from three specialized AI architectures: MedGemma (medgemma-4b-it-Q8_0, LLM for context extraction), HyenaDNA (for genomic risk prediction), and PLAPT AI (Transformer using molecular embeddings for chemical affinity screening). This end-to-end data fusion transforms raw genomic sequences and unstructured text into actionable therapeutic candidates, providing a scalable solution for proactive pandemic preparedness.

Therefore, the main pipeline has the following parts:

- Viral sentinel layer: NCBI Nucleotide API are used to scan for new viral complete genomes (DNA sequences, IDs, descriptions, host) from https://www.ncbi.nlm.nih.gov/nuccore. Why does the tool not use RefSeq, a curated, non-redundant reference database maintained by NCBI? The tool prioritizes access to the most recently submitted viral sequences and therefore does not rely on RefSeq (https://www.ncbi.nlm.nih.gov/refseq/), which requires additional time for expert curation. This design choice enables earlier detection of newly registered or emerging viruses, including potential zoonotic threats, without any delay introduced by the curation process.
- Contextual data enrichment with a LLM (Model 1): The pre-trained MedGemma (medgemma-4b-it-Q8_0, https://huggingface.co/lmstudio-community/medgemma-4b-it-GGUF) is used with LM Studio to correct and complete the host information for the viruses.
- Zoonotic score prediction with a new HyenaDNA-based topology (Model 2): We trained virsentai-v2-hyenadna-16k classifier (host as human and non-human) that will use as input the DNA sequences (base level) for new viruses and it will calculate the zoonotic score (probability of the virus to infect the humans or to have a host the humans). We fine-tunned hyenadna-medium-160k-seqlen-hf model (https://huggingface.co/LongSafari/hyenadna-medium-160k-seqlen-hf).
- Protein and drugs extraction layer: for the viruses with the predicted zoonotic score greater than 90%, the protein amino acid sequences from NCBI data are extracted (if exists). The FDA approved drugs are obtained from ChEMBL as SMILES formulas.
- Protein – drug interaction layer: the protein sequences of new viruses with high zoonotic predicted score and the SMILES of approved drugs are used as input for the PLAPT transformer-based pre-trained model (Model 3) in order to calculate the affinity interaction (neg_log10_affinity_M). This model is based on ProtBERT protein embeddings and ChemBERTa drug embeddings (https://github.com/Bindwell/PLAPT). Our group used PLAPT drug repurposing in two previous studies about therapeutic target in sepsis pathogenesis (Kirtan et al., 2025) and druggable cancer-driving proteins (López-Cortés et al., 2024).
- Data storage: the data from all the steps are stored into a SQLite database, TSV and CSV files.
- Data processing: python scripts are extracting zoonotic prediction and drug repurposing summaries as JSON / CSV files. Only viruses with zoonotic score >=90% and PLAPT negative log 10 affinity >= 10 (very strong interactions) are included. An additional filter for NCBI data that could not be real viruses is applied (using a list of keywords).
- Web tool update: the summaries are included into HTML pages such as zoonotic virus predictions and drug repurposing sections. These pages are plotting filtered data as tables, dynamic plots, and a virus – protein - drug interaction diagram.

### 2.1 VirSentAI model

We built our model, virsentai-v2-hyena-dna-16k, by fine-tuning the powerful HyenaDNA foundation model (https://huggingface.co/LongSafari/hyenadna-medium-160k-seqlen-hf), a transformer-like architecture pretrained on billions of DNA bases to learn the fundamental “language” of genomics (Nguyen et al., 2023; Consens et al., 2025). This is where the architecture’s true advantage lies. Its use of convolutions with learnable long-range filters—the so-called Hyena operators—allows it to process entire viral genomes up to 160,000 bases without resorting to the fragmenting or windowing techniques that can obscure critical information (effectively replacing the costly attention mechanism of traditional Transformer network for subquadratic scaling). The result is an engine uniquely capable of capturing the subtle, distributed sequence signals, from codon biases to complex structural motifs, which correlate with human tropism.

To specialize this powerful foundation for zoonotic surveillance, we undertook a meticulous fine-tuning process. The fuel for this was a curated training corpus of 31,728 complete viral genomes, drawn from three cornerstone repositories: the National Center for Biotechnology Information (NCBI; Sayers et al., 2024), VirusHostDB (Mihara et al., 2016), and the Virus Pathogen Resource (VIPR), which has been integrated into the BV-BRC (Olson et al., 2023). A critical step in this process was ensuring a strict balance between human and non-human host labels—a measure essential for mitigating classification bias. During training, each single-nucleotide tokenized sequence was fed into the model, which learned to predict a continuous human-infectivity score represented as a probability from 0 to 1.

The model was trained over 15 epochs using 16-bit mixed-precision and the AdamW optimizer. To manage the significant memory demands on a single 24GB NVIDIA GPU, we employed a small batch size of two, augmented by eight steps of gradient accumulation. This technique effectively simulates a larger batch size, stabilizing the learning process without requiring additional hardware. This entire 150-hour regimen was carefully calibrated to maximize performance within the very real constraints of typical academic research infrastructure.

### 2.2 Web Server Architecture

The backend logic of VirSentAI was implemented in Python, leveraging powerful libraries such as PyTorch for deep learning inference. The core function of the backend is to run the automated scanning and prediction pipeline. A lightweight SQLite database is used for storing prediction results, virus metadata, and summary statistics. This choice supports the self-contained and easily deployable nature of the platform.

The frontend user interface is built with standard web technologies, including HTML, CSS and JavaScript, to ensure compatibility across all modern browsers. The main page includes the “Top Predicted Threat” virus with its host taxonomy and probability, a table of the 10 most-zoonotic viruses among recent inputs, and dynamic chart as a time-series of predicted zoonotic viruses. All virus IDs are linked to the details from the NCBI database. The system is designed for ease of maintenance: new sequences are queried via NCBI API, HyenaDNA inference is fast on modern GPUs, and results are updated in real time on the main dashboard. **Figure S1** is presenting the main interface of the VirSentAI platform with zoonotic virus prediction scores.

**Figure S2** shows the PLAPT drug repurposing section with statistics about interactions between the viral proteins and approved drugs (PLAPT negative log 10 affinity >= 10 for very strong interactions). You can filter the interactions using virus ID, protein ID, drug ID and the affinity range. The drug repurposing section shows:

- Summaries such as the total number of interactions, the average affinity, the number of viruses, drugs and proteins in these results.
- A dynamic interaction networks with nodes as viruses, proteins and drugs.
- A virus-drug interactions heatmap.
- A bar plot for binding affinity distribution.
- A bar plot with the top high-affinity interactions.
- A table with the interaction data used in the previous statistics and graphical visualizations. All the virus IDs and protein IDs are linked to the NCBI database for details. The drug IDs are linked to the ChEMBL database details.

In line with our commitment to open science, the entire source code is available on GitHub at https://github.com/muntisa/virsentai and the platform is publicly accessible at https://muntisa.github.io/virsentai.

## 3. Results

The efficacy of a viral host prediction model is ultimately defined not just by its quantitative metrics, but also by its ability to distinguish a faint zoonotic signal from the overwhelming noise of viral diversity. VirSentAI demonstrates strong performance, achieving an area under the receiver operating characteristic curve (AUROC) of 0.9496 and an overall accuracy of 0.8724. These figures position it competitively among leading models in the domain, including specialized predictors that report AUROC scores in the 0.95–0.99 range (e.g., Brierley et al., 2024; Hu et al., 2025).

The progression of the field has moved from highly specialized tools toward more generalized models (**Table 1**). Early models offered high precision within narrow viral families but lacked the flexibility to generalize across a broader pathogenic landscape. The subsequent wave of alignment-free classifiers and recurrent neural networks expanded this horizon, but often at the cost of losing long-range sequence context. More recently, massive foundation models like Evo 2 signal a conceptual leap toward large-scale genomic learning, although their exclusion of pathogenic viruses from training data currently limits their direct applicability for host prediction.

**Table 1.**
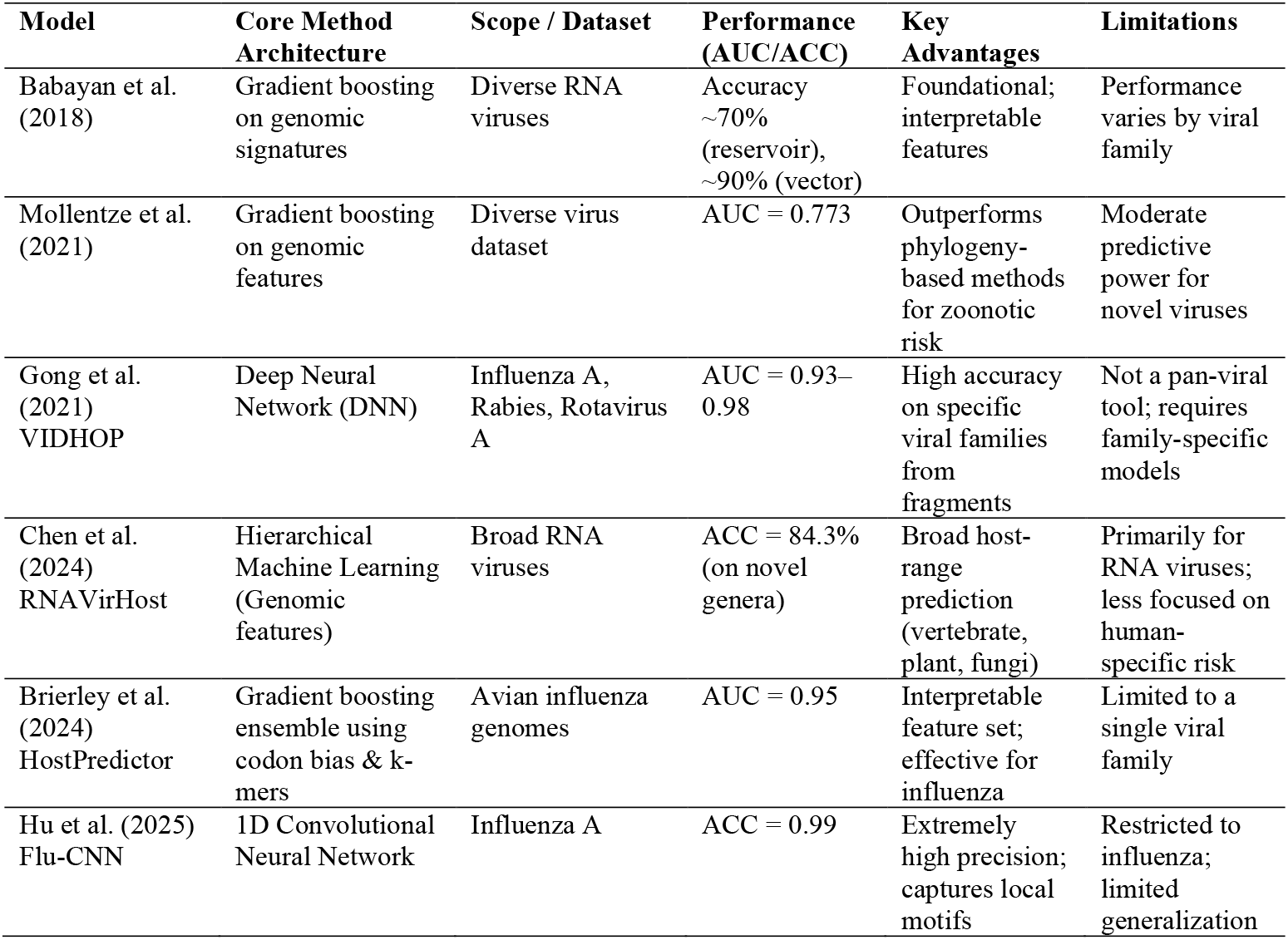
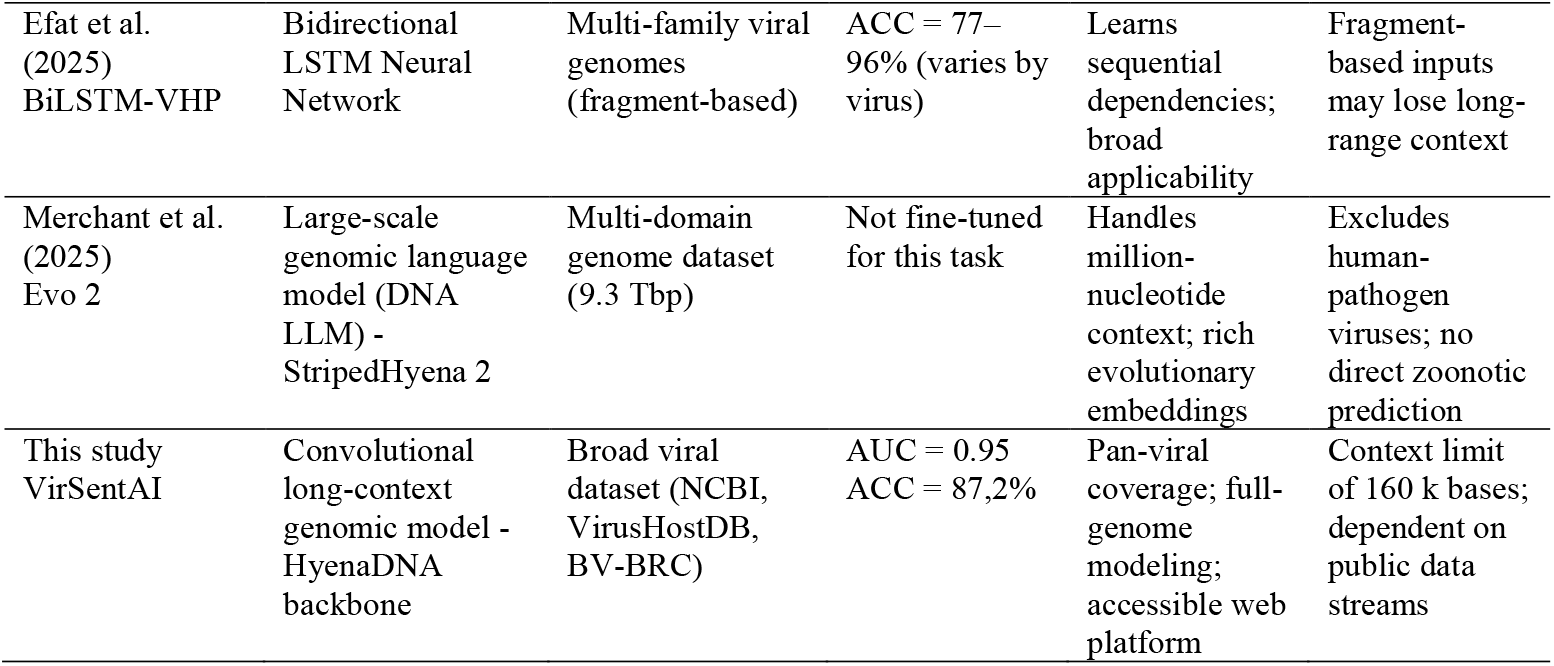
Comparison of representative models for viral host prediction.

VirSentAI was designed to bridge the gap between narrow specialists and broad, unspecialized foundation models. Its core architectural advantage lies in its ability to process entire viral genomes as single, continuous sequences at single-nucleotide resolution. This allows the model to capture not only local mutational signatures but also the long-range dependencies across the genome that are often crucial for host adaptation—context that is frequently lost in methods relying on fragmented or feature-engineered data.

This architectural choice carries significant practical implications. The HyenaDNA framework is computationally efficient: VirSentAI is a compact model of approximately 120 million parameters, trainable in under one week on a single 24GB GPU. This efficiency contrasts sharply with the resource-intensive demands of billion-parameter models and is a key factor in its viability as an operational tool rather than a purely academic one. The resulting platform integrates scientific rigor with real-world usability. Its data pipeline, directly linked to NCBI repositories, robustly handles the ingestion and preprocessing of large-scale genomic data. Up to current date, 16,060 viruses have been scanned (new and with unknown host) and 33 have been predicted to have zoonotic score >= 90%.

The database contains 29,625 viral protein – drug PLAPT interaction affinities >= 8.0 (on November 24, 2025) based on predicted zoonotic viruses with probability >= 80%. The data is filtered for the Web interface statistics by using cutoff for zoonotic score of 90% and PLAPT affinities of 10.0. The top 10 zoonotic viruses predicted by VirSentAI are presented in **Table 2**.

**Table 2.** Top 10 zoonotic viruses predicted by VirSentAI.

| Virus ID | Virus Name | Virus Host | Probability |
| --- | --- | --- | --- |
| NC_006502.1 | Isavirus salaris | Salmon | 98.15% |
| NC_043126.1 | Longquan virus | Bats, insectivores, rodents | 98.00% |
| NC_043363.1 | Choristoneura fumiferana entomopoxvirus | Choristoneura fumiferana | 97.90% |
| NC_074760.1 | Heterocapsa circularisquama DNA virus 01 | Heterocapsa circularisquama | 97.68% |
| NC_001753.1 | Clover yellow mosaic virus, complete genome | Unknown | 97.63% |
| NC_002205.1 | Influenza B virus (B/Lee/1940) | Unknown | 97.58% |
| NC_007750.1 | Peruvian horse sickness virus segment 3, complete genome | Vertebrates, mosquitoes | 97.40% |
| OM869523.1 | Sigmofec virus UA08Rod_6044, complete genome | Cricetidae, Heteromyidae | 97.00% |
| NC_074761.1 | Heterocapsa circularisquama DNA virus 01 | Heterocapsa circularisquama, Unknown | 96.98% |
| NC_004053.1 | Ophiostoma mitovirus 5, complete genome | Ophiostoma novo-ulmi | 96.89% |

The top PLAPT interactions between the approved drugs and the zoonotic virus proteins are presented in **Table 3**.

**Table 3.**
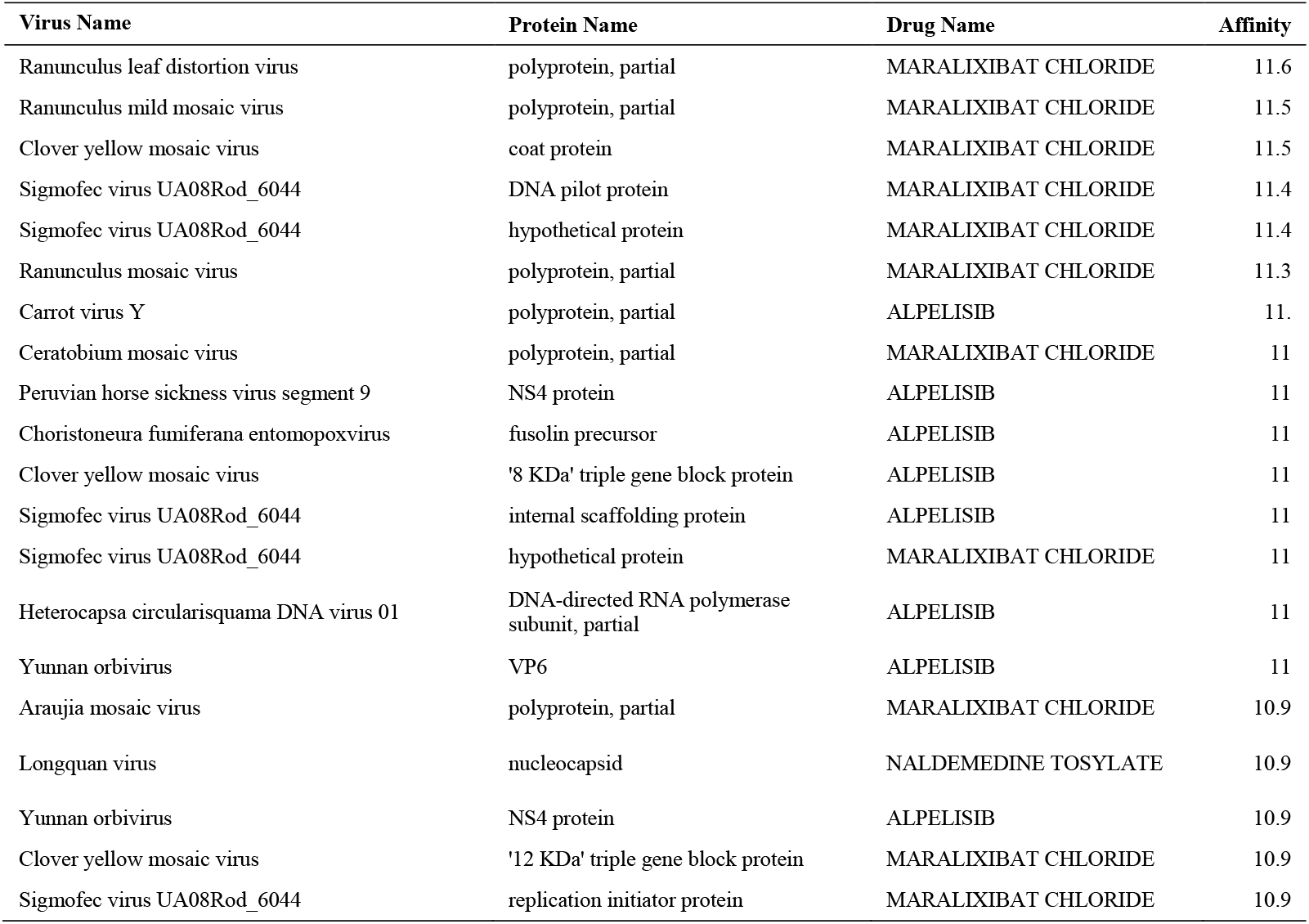
Top drugs that could interact with zoonotic virus proteins.

## 4. Discussion

VirSentAI is not without its limitations. The current 160k-token context window can be restrictive for the largest known viral genomes. Furthermore, the rapid convergence of the model during training suggests that extended training epochs could yield further performance improvements. The current implementation also relies solely on NCBI data. The PLAPT interaction model may introduce inaccuracies into the resulting drug repurposing predictions. The large language model employed to enhance virus metadata relies exclusively on NCBI data, which may limit its coverage and lead to failure in cases where information is absent unless external web resources are utilized. These aspects are not fundamental flaws but rather represent clear avenues for future refinement and development.

The application of a HyenaDNA-based AI model for predicting human host potential from complete viral genomes yielded a wide spectrum of results. This analysis reveals the model’s profound capabilities in genomic pattern recognition while simultaneously highlighting its critical deficiencies in biological and ecological reasoning. We have categorized the predictions to delineate the model’s successes, nuanced outputs, and theoretical failures.

### 4.1 Filtered zoonotic predictions

We first present the high-confidence zoonotic predictions corresponding to the data presented in **Table 2**, which reflects the current web results filtered for a prediction cutoff of 90% and excluding non-viral sequences. The model demonstrated exceptional accuracy in identifying viruses that are well-established as human pathogens or belong to high-risk zoonotic reservoirs, effectively acting as a high-throughput genomic classifier. The top predictions generated by the Autonomous VirSentAI Agent highlight its capability to identify potential cross-species transmission risks, often-flagging non-mammalian viruses with high genomic similarity to established human pathogens. The high prediction probabilities (96.89% or greater) across this diverse set underscore the robustness of the fine-tuned HyenaDNA topology in recognizing sequence patterns associated with human host adaptation, even when the known host is non-human or currently non-zoonotic.

These predictions validate the model’s core utility by identifying established human pathogens or viruses from high-risk zoonotic reservoirs:

- NC_043126.1 - Longquan virus (98.00%): This is a most compelling success. It validates the model’s ability to flag High-Consequence Zoonotic Viruses from classic mammalian reservoirs (Bats, insectivores, and rodents), which are associated with diseases like HFRS (Hantavirus Family). This confirms the model’s capacity to identify genuine emergent threats based on genomic signatures alone.
- NC_002205.1 - Influenza B virus (B/Lee/1940) (97.58%): This result is a crucial positive control, confirming the model’s ability to accurately detect the distinct genomic signature of a highly adapted human pathogen, even when the specific host field in the database is “Unknown.”
- NC_007750.1 - Peruvian horse sickness virus segment 3 (97.40%): As an arthropod-borne virus (*Orbivirus* family) infecting vertebrates, its high probability reinforces the model’s accurate placement of the virus into the broad “vertebrate virus” category, which is essential for identifying potential zoonotic agents transmitted by vectors like mosquitoes.
- OM869523.1 - Sigmofec virus UA08Rod_6044 (97.00%): This virus circulates in rodents (*Cricetidae* and *Heteromyidae*). This result confirms the model’s successful identification of high-risk signatures associated with critical mammalian reservoir hosts for numerous zoonotic pathogens, including hantaviruses and arenaviruses.
- NC_006502.1 - Isavirus salaris (98.15%): The strong prediction for this virus, traditionally restricted to Salmon, is significant because ISAV belongs to the *Orthomyxoviridae* family (like Influenza). The model suggests that the genomic composition of ISAV shares structural or functional patterns highly analogous to the general human host signature learned from its Influenza relatives, signaling a need for continued surveillance given the family’s known zoonotic potential.

These predictions highlight the core limitation of a purely sequence-based model - the inability to integrate biological and ecological barriers:

- NC_043363.1 - Choristoneura fumiferana entomopoxvirus (97.90%): The strong prediction for this insect-specific virus indicates its genomic signature shares key structural patterns with human host fingerprints. This identifies an unexpected genomic overlap but represents a biologically untenable prediction. The entire replication machinery of entomopoxviruses is adapted for insect cellular biology, making a cross-kingdom jump to vertebrates highly implausible.
- NC_001753.1 - Clover yellow mosaic virus (97.63%): The prediction that this plant virus could have a human host is a categorical error. The fundamental barriers (cellular receptors, replication machinery, and optimal temperature ranges) prevent such an infection.
- NC_004053.1 - Ophiostoma mitovirus 5 (96.89%): The strong prediction for this fungal virus is a categorical error. This exposes the core limitation where the HyenaDNA topology recognizes a highly conserved RNA sequence motif that randomly matches a genomic feature common to certain human RNA viruses, failing to account for the kingdom-level biological barriers.
- NC_074760.1 / NC_074761.1 - Heterocapsa circularisquama DNA virus 01 (97.68% / 96.98%): These predictions relate to a virus that infects marine algae. They represent a novel category of implausible predictions akin to protozoan viruses. The biological chasm between an algal host and a human cell makes a host jump not supported by any known virological principles.

### 4.2 Extended zoonotic predictions

The following analysis presents an extended view of results obtained with a lower cutoff of 80% and without the use of host filtering with rules and LLM. This unfiltered data includes many viruses where the host information was initially listed as non-human or unknown, demonstrating the comprehensive initial screening capability before filtering is applied.

The model demonstrated strong accuracy in classifying viruses not included in the training datasets but labelled with unknown hosts. These successes confirm the model’s ability to process complex genomic data and identify patterns consistent with known human pathogens. These are pandemic and epidemic human viruses:

- The most striking success was the correct classification of numerous isolates of Severe acute respiratory syndrome coronavirus 2 (SARS-CoV-2, OZ301499.1 and OZ302098.1, 87.62%) and the milder Influenza C virus (NC_006307.2, 84.85%).
- In addition, the models identified human rotaviruses such as Norovirus Hu/GII-4/Sakai2/2006/JP (AB447448.1) with a 97.63% probability score.
- Similarly, it correctly tagged multiple strains of human rotavirus (e.g., NC_021548.1 – 95.79%), a globally significant cause of childhood gastroenteritis.

Critically, the model correctly flagged viruses of significant zoonotic concern (high-consequence zoonotic viruses). The identification of Influenza A virus (NC_007362.1, 80.62%), specifically the A/goose/Guangdong/1/1996(H5N1) strain, is particularly noteworthy. This is the progenitor of the highly pathogenic H5N1 avian influenza, which has demonstrated its capacity for deadly, albeit inefficient, transmission to humans. Recognizing the latent threat in a non-human virus genome is a significant achievement and points to the real-world utility of this approach for pandemic preparedness.

These nuanced predictions involve vertebrate viruses with unconfirmed or low zoonotic potential (arboviruses and other mammalian viruses). The model correctly identifies them as belonging to vertebrate-associated viral families but appears to overstate their direct human host potential:

- The model flagged Banna virus (NC_004201.1, 86.39%). Both this virus and Peruvian horse sickness virus (97.40%, also in **Table 2**) are arthropod-borne viruses (arboviruses) that infect mammals. While some arboviruses are significant human pathogens (e.g., Dengue virus), the zoonotic potential of others, like Banna virus, is considered low or not well established.
- The model also identified Cottontail rabbit papillomavirus (NC_001541.1, 80.72%). While related to human papillomaviruses (HPVs), there is no evidence that this specific virus can cross the species barrier to infect humans. In these cases, the model correctly places the virus in a broad “vertebrate virus” category based on genomic features but lacks the fine-grained specificity to distinguish between a rabbit, horse, or human host.

The model’s inability to comprehend ecological relationships leads to contextually misleading predictions:

- The model consistently identifies bacteriophages commonly found in human microbiome samples as having a human host. An example is Bajarodmic virus 902 (PQ007151.1), which scored a remarkably high 98.46% probability. The genomic input is derived from a human environment, but the model misattributes the host, confusing a virus *in* humans (infecting our bacteria) with a virus *of* humans.

The model’s most significant failures occur when analyzing viruses from hosts that are evolutionarily distant from vertebrates (biologically implausible predictions, crossing kingdom-level barriers):

- Insect Viruses: Bracoviriform inaniti (NC_043270.1, 98.83%, “Top Predicted Zoonotic Threat”) and Daphnis nerii cypovirus (NC_040445.1, 94.62%). These are obligate pathogens of insects, adapted exclusively for insect cellular biology, making infection of a vertebrate biologically untenable.
- Viruses of non-mammalian vertebrates: The prediction for Iguanid herpesvirus 2 (NC_043063.1, 84.49%) is a prime example of the model stretching phylogenetic boundaries. Reptilian herpesviruses are adapted to cold-blooded hosts, and substantial barriers (including thermal incompatibility) prevent such a cross-class jump.
- Plant and Fungal Viruses: Rice gall dwarf virus (NC_009245.1, 85.73%). The prediction that this plant virus could have a human host is a categorical error due to absolute cellular and biological barriers.
- Protozoan Viruses: Paramecium bursaria Chlorella virus NC1A (NC_043234.1, 86.25%) and Giardia lamblia virus (NC_040632.1, 87.15%). These involve viruses that infect single-celled eukaryotes. The biological chasm between these hosts and human cells is immense, and the prediction of a host jump is not supported by any known virological principles.

### 4.3 *In silico* automatic drug repurposing for zoonotic viruses

The final stage of the Autonomous VirSentAI Agent workflow involves the application of the PLAPT AI model to perform a large-scale in silico drug repurposing analysis. This computational screening identifies potential binding interactions between the extracted proteins from predicted zoonotic viruses and a curated library of approved or clinically advanced small molecules (**Table 3**). This step transforms the initial genomic prediction (host risk) into actionable therapeutic intelligence (drug candidates).

The affinity scores in **Table 3** represent the predicted binding strength (affinity), often expressed as a p*K*_*i*_ or equivalent metric, where higher values indicate stronger predicted binding. Two primary candidates dominate the analysis (Maralixibat Chloride and Alpelisib), targeting a diverse range of viral proteins:

- Maralixibat Chloride (CHEMBL17879) is a drug currently approved for cholestasis in Alagille syndrome, and it is predicted to interact with the highest affinity (up to 11.6) across several viral polyproteins, particularly those derived from the Ranunculus and Clover yellow mosaic virus groups. These high-scoring interactions are predominantly with polyproteins, which are often cleaved into multiple functional enzymes and structural components critical for viral replication. The strong signal across multiple virus polyproteins suggests a potential inhibitory mechanism targeting a highly conserved motif present within the proteolytic machinery or replication complex of these viruses, irrespective of the specific host (plant, rodent, etc.). Given that Maralixibat is a bile acid transporter inhibitor, its predicted activity against viral proteins is likely an off-target effect captured by the structure-based PLAPT AI model. While these viruses are flagged as biologically implausible zoonotic risks in the initial HyenaDNA screening, the successful identification of a high-affinity small molecule provides novel structural templates for pan-viral drug design, provided the interaction can be validated experimentally.
- Alpelisib (CHEMBL2396661) is an inhibitor of the phosphoinositide 3-kinase (PI3K) pathway used in oncology, and it demonstrates strong predicted affinity (up to 11.0) against several key regulatory and structural proteins, including:
  ∘ NC_007753.1 - Peruvian horse sickness virus NS4 protein: This virus is a confirmed arbovirus threat. The NS4 protein is non-structural and often critical for host immune evasion and replication. Targeting a regulatory protein of a vertebrate pathogen with an established kinase inhibitor offers a mechanistically plausible path for repurposing, as many mammalian viruses exploit host kinase signaling pathways.
  ∘ NC_043363.1 - Choristoneura fumiferana entomopoxvirus fusolin precursor: Although the virus is insect-specific, the predicted high binding to the fusolin precursor (a protein involved in the dissemination and stability of the virus) offers a structural hypothesis for inhibition if this motif is conserved across related poxviruses with true zoonotic potential (e.g., orthopoxviruses).
  ∘ OM869523.1 - Sigmofec virus (Rodent reservoir) is an internal scaffolding protein: This is another highly relevant prediction against a virus from a known high-risk mammalian reservoir. The internal scaffolding protein is essential for virion assembly. Inhibiting this structural step with Alpelisib warrants urgent experimental validation, as disrupting virus assembly is a proven antiviral strategy.
- Naldemedine Tosylate (CHEMBL3039508) is an opioid antagonist, presents a high-affinity interaction (10.9) with the Longquan virus nucleocapsid protein (YP_009664869.1). The Longquan virus is a high-consequence zoonotic Hantavirus. The nucleocapsid protein is vital for both genome encapsidation and transcription, making it an excellent and highly conserved antiviral target. This prediction against a Tier 1 zoonotic threat from the HyenaDNA screen represents the most clinically relevant repurposing hypothesis presented in **Table 3**. The predicted inhibition of this critical viral function by a non-antiviral drug requires immediate investigation as a promising therapeutic strategy.

The results demonstrate the core value of the VirSentAI Agent’s dual-model approach: transforming genomic prediction into structural/therapeutic hypothesis generation. The utility of the PLAPT AI model lies in its ability to quickly identify privileged scaffolds (like those in Maralixibat and Alpelisib) that can inhibit viral processes through unexpected off-target effects. However, the *in silico* nature of the predictions necessitates caution:

- Biological implausibility: The strong predictions against viruses previously flagged as biologically implausible human pathogens (plant and fungal viruses) likely stem from conserved protein domains. While these interactions are structurally valid, they are not biologically actionable unless the conserved protein domain is also found in a bona fide human pathogen.
- Affinity vs. efficacy: High predicted binding affinity does not guarantee biological efficacy or favorable pharmacokinetics within a human host. The proposed drug-protein interactions must be validated through in vitro assays (e.g., *IC*_*50*_ determination) and subsequent in vivo models to confirm their antiviral effect.

In conclusion, the VirSentAI drug repurposing module has delivered several highly specific, novel therapeutic hypotheses against critical vertebrate and vector-borne viral threats, most notably Naldemedine Tosylate against the Longquan virus. These hypotheses define the immediate research priority for pre-clinical validation.

These results confirms that a sophisticated genomic model like HyenaDNA functions as a powerful but context-blind pattern recognizer. Its performance is exceptional for identifying established human pathogens and high-risk zoonotic agents like H5N1, suggesting immense potential as a primary screening tool for prioritizing threats from novel vertebrate viruses. However, its accuracy degrades significantly when assessing viruses from more distant hosts. The model consistently mistakes ecological presence for direct infection (bacteriophages) and is unable to comprehend the fundamental biological barriers that prevent viruses from jumping between different kingdoms of life. The future of AI in pandemic prediction lies not only in more powerful sequence analysis but also in the critical integration of biological, ecological, and structural data to provide the context that pure genomics cannot.

## 5. Perspectives on Future Work

Looking ahead, we are focused on transforming VirSentAI from a powerful prototype into a robust, high-availability surveillance platform. A key first step will be migrating to a dedicated Python-based web server and application, which will allow us to scale operations far beyond the current static GitHub page.

To broaden our detection scope, we plan to significantly extend the analysis window beyond the current 160 kb to effectively process the genomes of much larger viruses. On the data front, we aim to integrate additional genomic databases by leveraging their APIs and employing LLMs to rapidly scan and process incoming data. Crucially, we will integrate LLMs to summarize comprehensive data about predicted high-risk viruses from multiple sources, providing immediate, actionable insights. The model will improve with the integration of biological, ecological, and structural data to provide the context for pure genomics predictions.

Finally, we intend to move beyond simple risk scoring by integrating data on viral protein interactions with human proteins to better predict the consequences of infection. This will be coupled with an integration of drug and metabolite interaction data with the viral proteins, offering a fast-track route for drug repurposing against emerging threats.

Naturally, we will also continue to iterate on and improve all underlying prediction models to enhance accuracy and reliability across the board. The ideal viral sentry system it will be connected to real time PCR to detect viruses (Watzinger et al., 2006) with virtual detection of possible proteins from the new viral sequences in order to detect real-time viruses that could infect humans.

## 6. Conclusion

In pushing virus discovery forward, we have created something more than just another predictive model. VirSentAI represents a deliberate shift towards a proactive, rather than reactive, stance in genomic surveillance. The core idea was to build an intelligent, end-to-end platform capable of scanning the near-infinite horizon of genomic data for brewing threats, and—crucially—to make it profoundly accessible. Pandemics, after all, do not wait for command-line expertise.

We have engineered VirSentAI to scout for zoonotic risks directly from the raw dialect of viral DNA, harnessing the power of a long-range genomic model like HyenaDNA, which excels at interpreting vast, complex sequences. This is not just an incremental improvement. It is about fundamentally changing who gets to ask the questions. By wrapping this analytical prowess within a fully automated, open-source web service, we bridge a critical gap, empowering researchers and public health officials globally, regardless of their computational background. The entire platform, from its underlying code to a live dashboard, is freely available, a deliberate choice to foster global vigilance.

Of course, the platform has its limitations. No tool is a panacea. The resolution for pinpointing specific hosts can get fuzzy in certain edge cases, and the inherent biases lurking within training datasets remain a persistent shadow we must continually work to mitigate. Machine learning models, however powerful, are shaped by the data they learn from, a challenge across the field of viral prediction.

Still, the impact is a tangible step forward: bolstering surveillance, helping to spot faint patterns in the overwhelming noise of metagenomics data, and shining a light on potential epidemiological blind spots. The applications begin to sprawl from here. One could imagine synthetic virology labs using the platform to vet designs or API extensions hooking directly into global alert systems. This is where the method starts to show its true potential. By creating a tool that is both powerful and open, VirSentAI nudges the global health community toward a more democratized and anticipatory posture in the unending race against the next pandemic threat.

## Data and Code Availability

- Web Server: https://muntisa.github.io/virsentai
- Source Code: https://github.com/muntisa/virsentai
- Release v1.1.0: https://doi.org/10.5281/zenodo.17445222

Note: All data, analysis scripts, and details about the models supporting the findings of this study will be made publicly available in a permanent repository upon publication.

## Acknowledgments

The CITIC is funded by the Xunta de Galicia through the collaboration agreement between the Consellería de Cultura, Educación, Formación Profesional e Universidades, and the Galician universities for the strengthening of research centers in the Galician University System (CIGUS). This work was supported by the Consolidation and Structuring of Competitive Research Units (ED431C 2022/46), GRC funded by Xunta de Galicia endowed with EU FEDER funds.

