## Supplementary material for "Viral Sentry AI (VirSentAI) - Automated Zoonotic Surveillance & Drug Repurposing Agent": Figure S1 and Figure S2

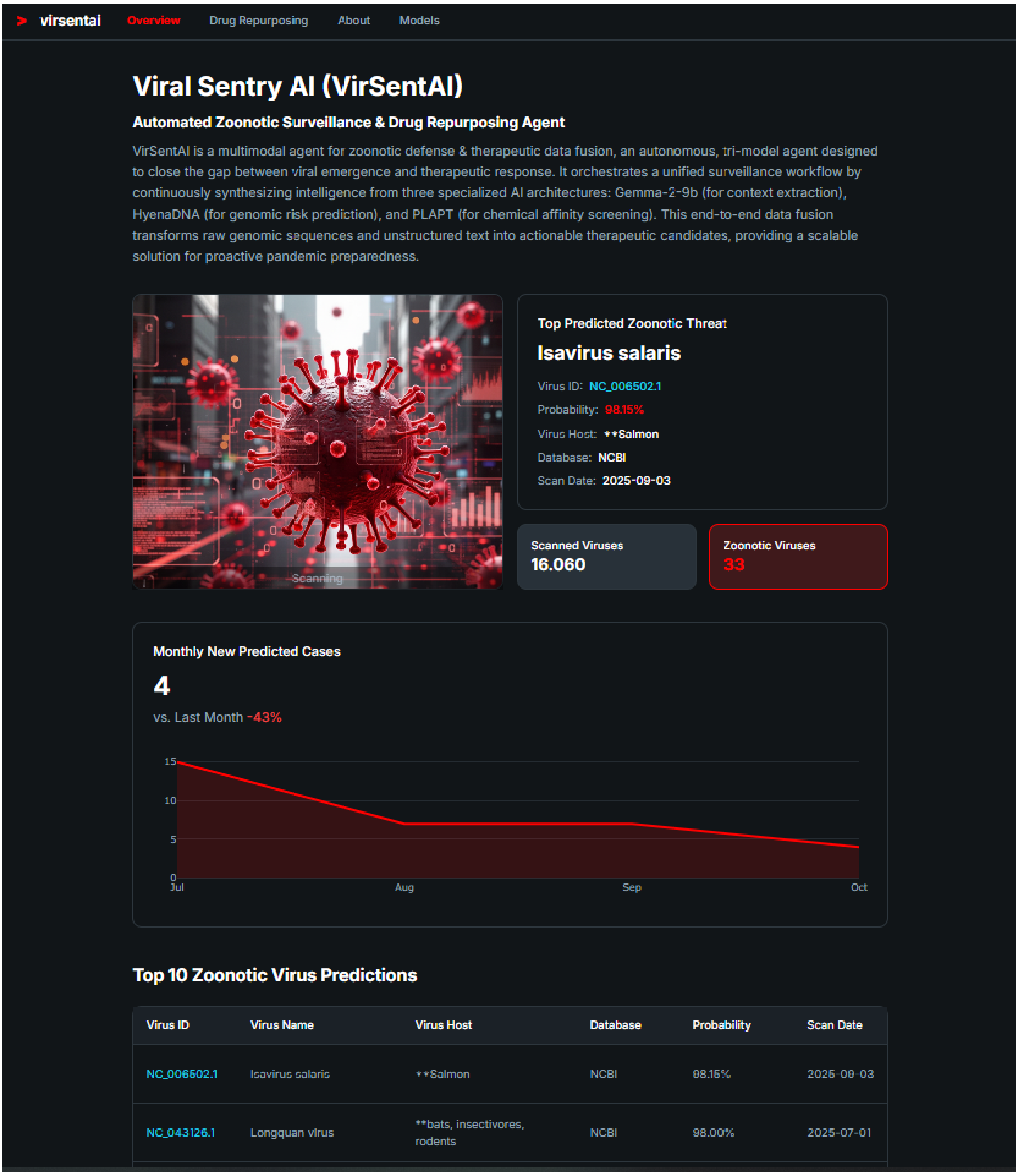


**Figure S1**: Screenshot of the VirSentAI main page: zoonotic virus predictions (24/12/2025).

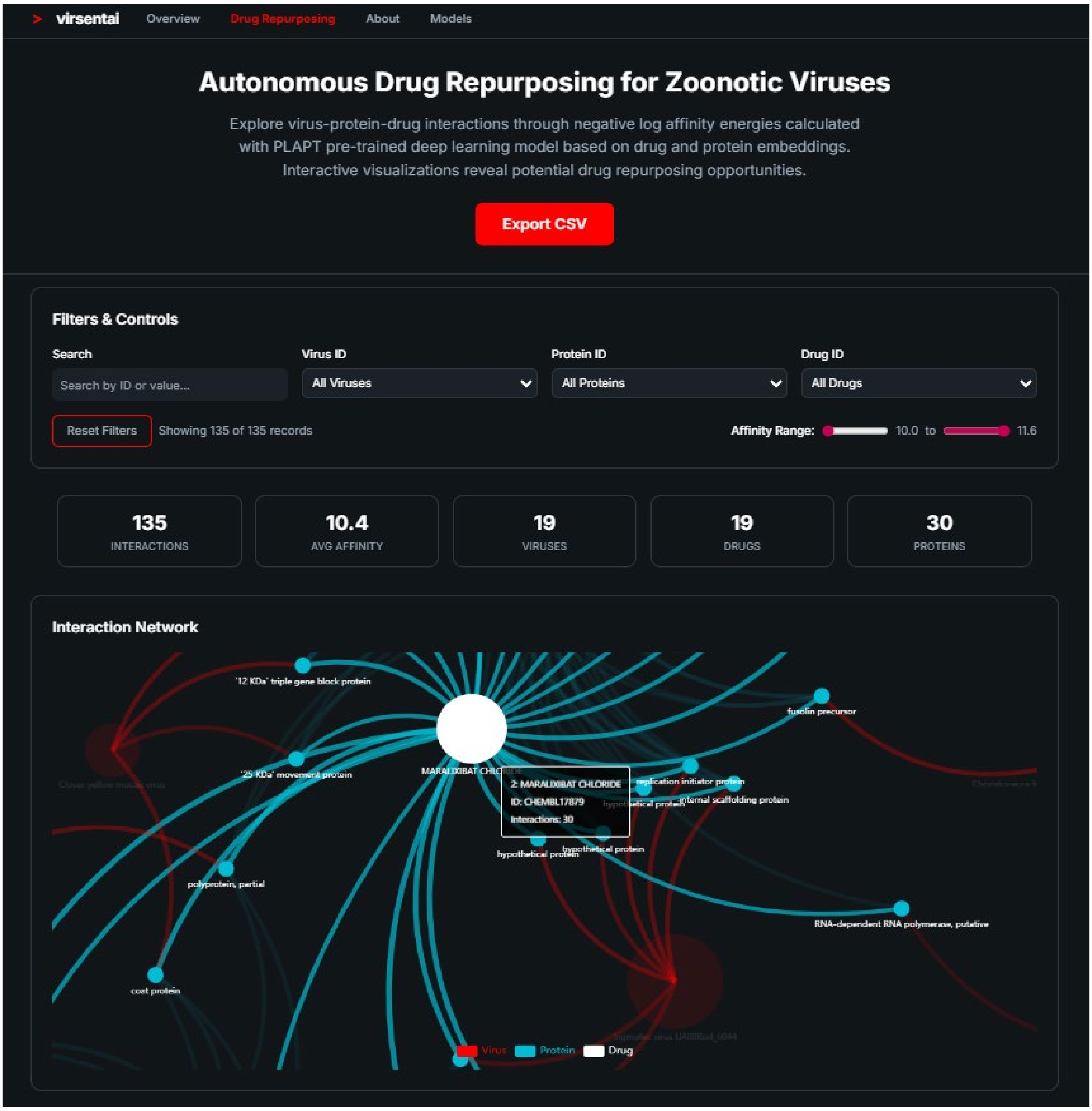


**Figure S2**: Screenshot of the VirSentAI drug repurposing section (24/12/2025).
